# Maternal BMI at the start of pregnancy and offspring epigenome-wide DNA methylation: Findings from the Pregnancy and Childhood Epigenetics (PACE) consortium

**DOI:** 10.1101/125492

**Authors:** Gemma C Sharp, Lucas A Salas, Claire Monnereau, Catherine Allard, Paul Yousefi, Todd M Everson, Jon Bohlin, Zongli Xu, Rae-Chi Huang, Sarah E Reese, Cheng-Jian Xu, Nour Baïz, Cathrine Hoyo, Golareh Agha, Ritu Roy, John W Holloway, Akram Ghantous, Simon Kebede Merid, Kelly M Bakulski, Leanne K Küpers, Hongmei Zhang, Rebecca C Richmond, Christian M Page, Liesbeth Duijts, Rolv T Lie, Phillip E Melton, Judith M Vonk, Ellen A Nohr, CharLynda Williams-DeVane, Karen Huen, Sheryl L Rifas-Shiman, Carlos Ruiz-Arenas, Semira Gonseth, Faisal I Rezwan, Zdenko Herceg, Sandra Ekström, Lisa Croen, Fahimeh Falahi, Patrice Perron, Margaret R Karagas, Bilal Mohammed Quraishi, Matthew Suderman, Maria C Magnus, Vincent WV Jaddoe, Jack A Taylor, Denise Anderson, Shanshan Zhao, Henriette A Smit, Michele J Josey, Asa Bradman, Andrea A Baccarelli, Mariona Bustamante, Siri E Håberg, Göran Pershagen, Irva Hertz-Picciotto, Craig Newschaffer, Eva Corpeleijn, Luigi Bouchard, Debbie A Lawlor, Rachel L Maguire, Lisa F Barcellos, George Davey Smith, Brenda Eskenazi, Wilfried Karmaus, Carmen J Marsit, Marie-France Hivert, Harold Snieder, M Daniele Fallin, Erik Melén, Monica C Munthe-Kaas, Hasan Arshad, Joseph L Wiemels, Isabella Annesi-Maesano, Martine Vrijheid, Emily Oken, Nina Holland, Susan K Murphy, Thorkild IA Sørensen, Gerard H Koppelman, John P Newnham, Allen J Wilcox, Wenche Nystad, Stephanie J London, Janine F Felix, Caroline L Relton

## Abstract

Pre-pregnancy maternal obesity is associated with adverse offspring outcomes at birth and later in life. Individual studies have shown that epigenetic modifications such as DNA methylation could contribute.

Within the Pregnancy and Childhood Epigenetics (PACE) Consortium, we meta-analysed the association between pre-pregnancy maternal BMI and methylation at over 450,000 sites in newborn blood DNA, across 19 cohorts (9,340 mother-newborn pairs). We attempted to infer causality by comparing effects of maternal versus paternal BMI and incorporating genetic variation. In four additional cohorts (1,817 mother-child pairs), we meta-analysed the association between maternal BMI at the start of pregnancy and blood methylation in adolescents.

In newborns, maternal BMI was associated with small (<0.2% per BMI unit (1kg/m^2^), P<1.06*10^-7^) methylation variation at 9,044 sites throughout the genome. Adjustment for estimated cell proportions greatly attenuated the number of significant CpGs to 104, including 86 sites common to the unadjusted model. At 72/86 sites, the direction of association was the same in newborns and adolescents, suggesting persistence of signals. However, we found evidence for a causal intrauterine effect of maternal BMI on newborn methylation at just 8/86 sites.

In conclusion, this well-powered analysis identified robust associations between maternal adiposity and variations in newborn blood DNA methylation, but these small effects may be better explained by genetic or lifestyle factors than a causal intrauterine mechanism. This highlights the need for large-scale collaborative approaches and the application of causal inference techniques in epigenetic epidemiology.

## Introduction

Offspring of mothers with a high body mass index (BMI) at the start of pregnancy have a higher risk of obesity and obesity-related disorders in later life(1). Maternal obesity in pregnancy is also associated with other offspring outcomes, including neurodevelopmental and respiratory outcomes(2–5). These associations might be explained by shared mother-child genetic or postnatal environmental influences, or they could also reflect a causal intrauterine mechanism leading to early programming of adverse health in the offspring(6).

Disentangling the genetic and shared postnatal environmental effects from a causal intrauterine effect is difficult, but there are a number of causal inference approaches that may be useful(7). For example, some studies have used a negative control design whereby the association between maternal adiposity and offspring outcome is compared to the association between paternal adiposity and the same outcome. The key assumption of the negative control design is that both exposures share the same postnatal environmental and genetic confounders. A systematic review(8) of such studies, together with subsequent studies not included in the review(9–12), have found only limited support for specific effects of maternal adiposity on offspring adiposity beyond birth. To our knowledge, similar causal inference techniques have not yet been applied to study maternal effects of adiposity in pregnancy on other aspects of offspring health.

If there is a causal intrauterine effect of maternal adiposity on offspring health outcomes, the mechanism is unclear. Epigenetic modifications, such as DNA methylation, might partly mediate associations between maternal and offspring phenotypes by causing changes to gene expression that are mitotically heritable(6, 13–15). Differential DNA methylation has been reported when assessing offspring exposed *in utero* to extreme maternal undernutrition(16–19), maternal morbid obesity(20) and less extreme maternal underweight and maternal obesity(21), in comparison to those not exposed; yet weak or no evidence has been found for associations between continuous maternal BMI and offspring DNA methylation, whether globally(22, 23), at specific loci identified in array(21, 24, 25) or at candidate genes(26). However, individual studies were limited in sample size and thus underpowered to detect differential methylation. Meta-analysis of results from multiple individual cohorts increases sample size and power to detect differential methylation, but this approach has rarely been employed in the field of epigenetic epidemiology.

Comprising many birth cohorts from around the world, the Pregnancy and Childhood Epigenetics (PACE) Consortium^25^ was established to facilitate meta-analysis of epigenomewide studies relevant to maternal and childhood health and disease. In this PACE study, we meta-analysed harmonised cohort-specific epigenome-wide data on associations between maternal BMI at the start of pregnancy and DNA methylation in the blood of newborns. We then conducted further analyses (Figure 1) to explore whether these associations could be reproduced in adolescent samples and implemented causal inference methods to evaluate the potential confounding effects of shared environment and genetic variation.

**Figure 1.**
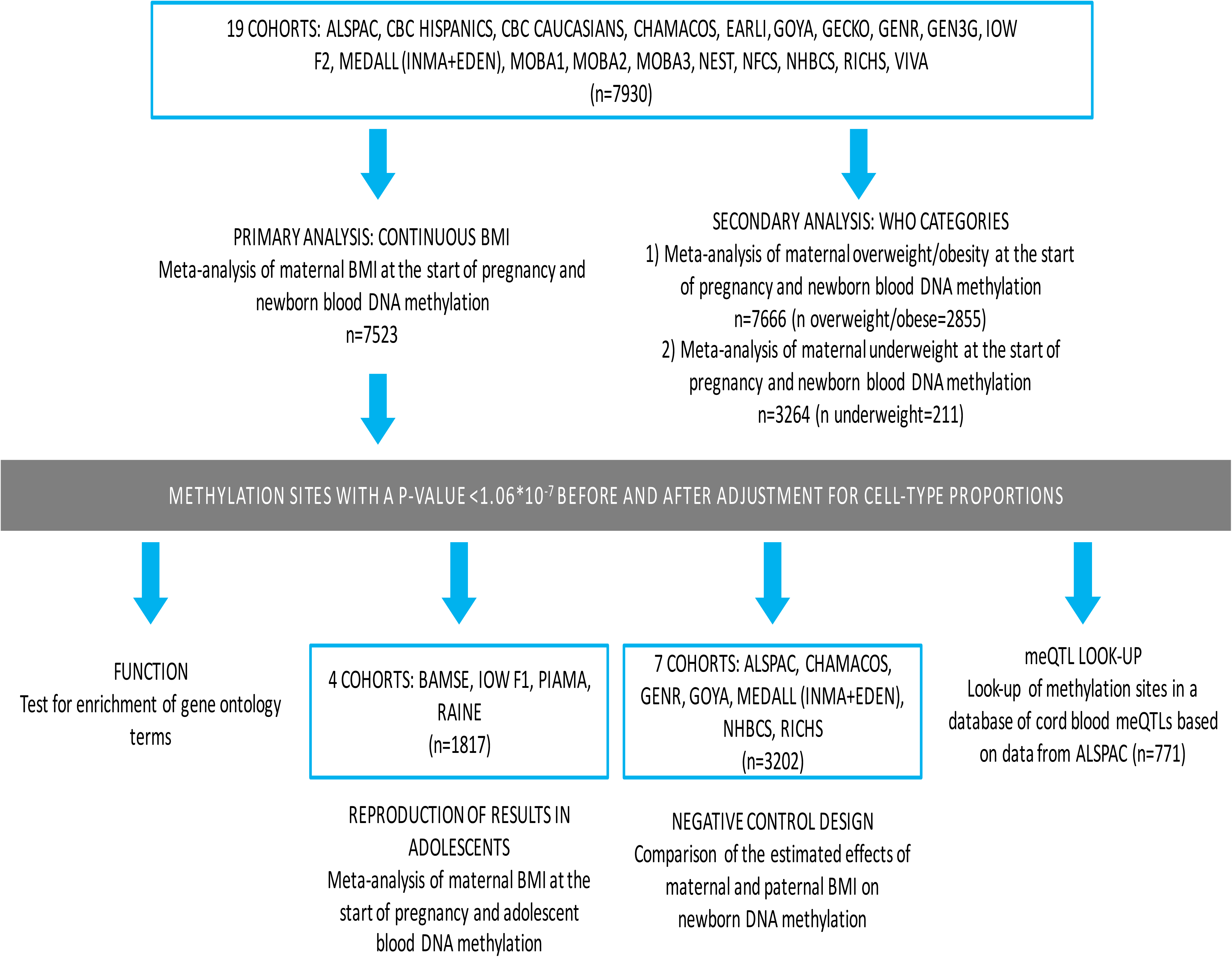
An overview of the study design

## Results

### Study characteristics

We meta-analysed results from 19 independent cohorts to test the association between maternal BMI at the start of pregnancy and epigenome-wide newborn blood DNA methylation. A summary of methods used by each cohort is provided in Supplementary File S1 Table S1, with a more detailed description in Supplementary File S2. Supplementary File S1 Table S2 lists sample sizes and summarises EWAS results for each cohort and meta-analysis. For our primary model, with continuous maternal BMI as the exposure, we analysed results from 7,523 mother-child pairs. The overall sample size-weighted mean maternal BMI was 24.4 kg/m^2^ (range of cohort-specific means: 22.8, 27.8). In secondary analyses, we examined World Health Organisation categories for maternal BMI, comparing normal weight women (n=4,834) to i) overweight or obese women combined (n=2,885 women, of whom 1,299 were obese) and ii) underweight women (n=211 women). The majority of participants were of European ancestry. Table 1 summarizes the characteristics of each cohort.

**Table 1.** Characteristics of each cohort included in the meta-analysis of the association between maternal pre-pregnancy BMI and offspring blood DNA methylation at birth. BMI is categorised according to WHO guidelines.

| COHORT | N IN<br>CONTINUOUS<br>BMI MODEL | MEAN MATERNAL<br>BMI (SD)<br>IN CONTINUOUS BMI<br>MODEL | MEAN MATERNAL<br>AGE (SD)<br>IN CONTINUOUS BMI<br>MODEL | TOTAL<br>N OBESE | TOTAL<br>N<br>OVERWEIGHT | TOTAL<br>N<br>UNDERWEIGHT | TOTAL<br>N<br>NORMAL<br>WEIGHT | ETHNICITY |
| --- | --- | --- | --- | --- | --- | --- | --- | --- |
| ALSPAC | 788 | 22.8 (3.6) | 29.7 (4.4) | 37 | 106 | 26 | 619 | European |
| CBC (Hispanic) | 132 | 24.2 (5.7) | 27.2 (5.7) | 15 | 27 | 11 | 79 | Hispanic |
| CBC (White) | 155 | 23.3 (3.9) | 32.0 (5.7) | 8 | 34 | 0* | 108 | European |
| CHAMACOS | 368 | 26.9 (5.1) | 25.3 (5.0) | 80 | 141 | 3* | 144 | Hispanic |
| EARLI | 211 | 27.8 (6.9) | 34.0 (4.7) | 69 | 51 | 3* | 88 | European/<br>Mixed |
| GECKO | 176 | 24.2 (3.9) | 30.4 (4.0) | 14 | 45 | 3* | 114 | European |
| GEN3G | 170 | 24.8 (5.6) | 28.0 (4.1) | 25 | 33 | 3* | 109 | European |
| Generation R | 875 | 24.5 (4.2) | 31.5 (4.2) | 90 | 202 | 13* | 570 | European |
| GOYA ** | 545 | 23.1 (3.2) | 29.5 (4.1) | 466 | 106 | 16 | 387 | European |
| IOW F2 | 53 | 27.7 (7.3) | 21.5 (1.4) | 19* | 11* | 0* | 23 | European |
| MEDALL<br>(INMA+EDEN) | 330 | 24.1 (5.1) | 30.6 (4.5) | 37 | 62 | 26 | 205 | European |
| MoBa1 | 1034 | 24.0 (4.6) | 29.9 (4.3) | 98 | 215 | 67 | 688 | European |
| MoBa2 | 647 | 24.2 (4.4) | 30.0 (4.5) | 72 | 136 | 18 | 431 | European |
| MoBa3 | 231 | 24.2 (4.3) | 29.6 (4.4) | 25 | 49 | 5* | 152 | European |
| NEST | 384 | 27.6 (8.9) | 28.8 (6.4) | 108 | 76 | 19* | 181 | Mixed |
| NFCS | 867 | 23.5 (4.1) | 29.1 (4.9) | 70 | 157 | 37 | 603 | European |
| NHBCS | 118 | 24.4 (4.2) | 31.0 (4.4) | 12 | 29 | 3* | 74 | European |
| RICHs | 96 | 25.8 (6.9) | 28.3 (5.5) | 21 | 21 | 10 | 44 | European |
| Project Viva | 343 | 24.3 (4.9) | 33.1 (4.5) | 41 | 77 | 10* | 215 | European |
| **Meta-analysis** | **7523** |  |  |  |  |  |  |  |

\*Included in the continuous BMI model, but excluded from the categorical analyses due to low sample sizes
\*\* A subset of the GOYA cohort (545) was included in the continuous BMI model. The entire cohort (975) was included in the binary BMI models.

### Maternal BMI at the start of pregnancy is associated with widespread but small differences in newborn blood DNA methylation

When treated as a continuous variable, maternal BMI at the start of pregnancy was associated with differential methylation in newborn blood at 9,044 sites (Supplementary File S1 Table S3) before and 104 sites (Supplementary File S1 Table S4) after adjustment for cell-counts (Bonferroni correction for 473,864 tests P<1.06*10^-7^); 86 sites were common to both models. Before adjustment for cell-counts, lambdas (*λ*), a measure of P-value inflation, were generally high and QQ plots showed inflation of P-values in most cohorts (Table 2, Supplementary File S1 Table S2, Supplementary File S3). Values for *λ* were closer to 1 for most cohorts after adjustment for estimated cell counts. In a meta-analysis of results from two of the larger cohorts, ALSPAC and Generation R (*λ*=1.60), *λ* was not substantially further reduced after removal of potential outliers using the Tukey method(27) (*λ*=1.58) or additional adjustment for 10 ancestry principal components (*λ*=1.67).

**Table 2.** Summary of cohort-specific and meta-analysis results for EWAS of continuous maternal pre-pregnancy BMI and newborn blood DNA methylation.

| COHORT | N | LAMBDA<br>(BEFORE<br>ADJUSTING<br>FOR CELLS) | BONFERRONI<br>HITS (BEFORE<br>ADJUSTING<br>FOR CELLS) | LAMBDA<br>(AFTER<br>ADJUSTING<br>FOR CELLS) | BONFERRONI<br>HITS (AFTER<br>ADJUSTING<br>FOR CELLS) |
| --- | --- | --- | --- | --- | --- |
| ALSPAC | 788 | 1.53 | 12 | 1.18 | 1 |
| CBC (Hispanic) | 132 | 1.05 | 12 | 0.96 | 7 |
| CBC2 (White) | 155 | 1.80 | 31 | 1.19 | 3 |
| CHAMACOS | 368 | 1.34 | 1 | 0.87 | 0 |
| EARLI | 211 | 0.88 | 0 | 0.89 | 2 |
| GECKO | 176 | 1.75 | 14 | 1.15 | 2 |
| GEN3G | 170 | 1.13 | 10 | 1.04 | 10 |
| GENR | 875 | 1.86 | 248 | 1.96 | 11 |
| GOYA | 545 | 1.87 | 2 | 1.01 | 1 |
| IOW F2 | 53 | 1.08 | 0 | 1.05 | 0 |
| MEDALL (INMA+EDEN) | 330 | 1.24 | 0 | 0.92 | 0 |
| MoBa1 | 1034 | 4.69 | 39 | 2.74 | 1 |
| MoBa2 | 647 | 2.70 | 8 | 2.76 | 14 |
| MoBa3 | 231 | 1.03 | 0 | 0.78 | 1 |
| NEST | 384 | 0.76 | 0 | 0.93 | 0 |
| NFCS | 867 | 0.95 | 0 | 0.98 | 0 |
| NHBCS | 118 | 1.02 | 2 | 1.17 | 4 |
| RICHs | 96 | 1.89 | 14 | 2.92 | 33 |
| VIVA | 343 | 1.27 | 8 | 1.49 | 7 |
| *FE Meta-analysis* | **7523** | **3.27** | **9044** | **2.41** | **104** |
| *RE Meta-analysis* |  |  | **1825** |  | **25** |

Sites associated with maternal BMI were spread over the genome and did not tend to be restricted to certain regions (Figure 2). Effect sizes were very small, with the median absolute effect at the genome-wide significant sites being a difference in methylation beta value of 0.0003 per one unit (kg/m^2^) increase in maternal BMI (i.e. a 0.03% absolute change, range: 0.15% decrease to 0.13% increase). At most of the Bonferroni-significant sites (8,899/9,044 and 96/104), higher maternal BMI was associated with lower newborn blood methylation.

**Figure 2.**
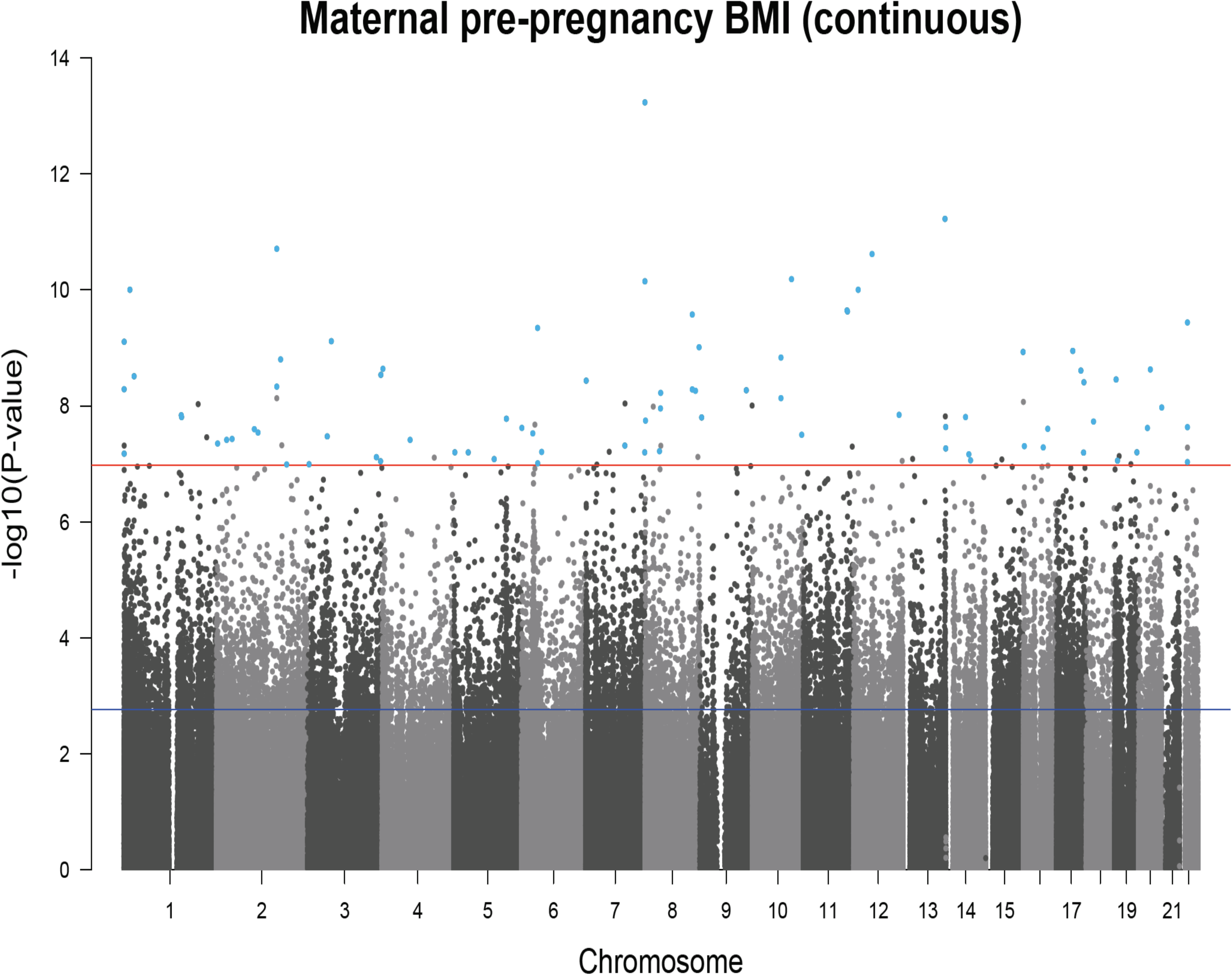
A Manhattan plot for the meta-analysis of associations between maternal pre-pregnancy BMI and offspring DNA methylation at birth after adjustment for maternal covariates and estimated cell counts. The red line shows the Bonferroni threshold for multiple testing. Methylation sites that surpassed the Bonferroni-correction threshold (P<1.06*10-7) before and after adjustment for estimated cell counts are highlighted in blue.

Results from the primary model, where the exposure was continuous BMI, were consistent with those from a binary comparison of maternal overweight/obesity (BMI>25) with normal weight (BMI 18.5 to 25): the Spearman’s coefficient for correlation between regression coefficients was 0.70. Maternal overweight/obesity was associated with differential newborn blood methylation at 4,037 sites (Supplementary File S1 Table S5) before and 159 sites (Supplementary File S1 Table S6) after cell-adjustment (P<1.06*10^-7^), compared with normal weight. The crossover between these 159 sites and the 104 identified with P<1.06*10^-7^ in the cell-adjusted continuous model was just 21/104, but 150/159 were associated with continuous BMI after correction for multiple testing at 159 sites (FDR-corrected P<0.05). The direction of effect for the binary comparison was consistent with that for the continuous exposure at all 159 sites. As expected, the magnitude of effect was larger when BMI was binary than when BMI was continuous, but the median effect at sites with P<1.06 *10^-7^ was still small (0.31% decrease in mean methylation beta value in the overweight/obese group compared to the normal weight group).

Eight sites (Supplementary File S1 Table S7) were associated with maternal underweight (BMI<18.5) compared to normal weight with P<1.06*10^-7^, but this analysis was likely underpowered given the small number of underweight women (n=211), and there was large inter-study heterogeneity in results (I^2^ median 62.3, range 0 to 91.3). Given these results, we did not explore the association between maternal underweight and offspring methylation any further.

### Adjusting for cellular heterogeneity greatly attenuates associations between maternal BMI and newborn blood DNA methylation

As mentioned above, adjusting for estimated cell proportions in newborn blood samples greatly reduced the number of sites associated with maternal BMI with P-values<1.06*10^-7^ (Figure 3). This reduction in signal was seen in all meta-analyses and most individual cohort analyses (Table 2). At all 9,044 sites associated with continuous maternal BMI, adjusting for cell counts shifted the effect size towards the null. The median relative change in estimate after adjustment was 52% and 9,007/9,044 sites attenuated by 10% or more. After adjustment, the precision of the estimates at 8,984/9,044 sites was increased (i.e. the standard error was reduced). Taken together, this suggests that much of the association between maternal BMI at the start of pregnancy and newborn DNA methylation is due to varying cell type proportions.

**Figure 3.**
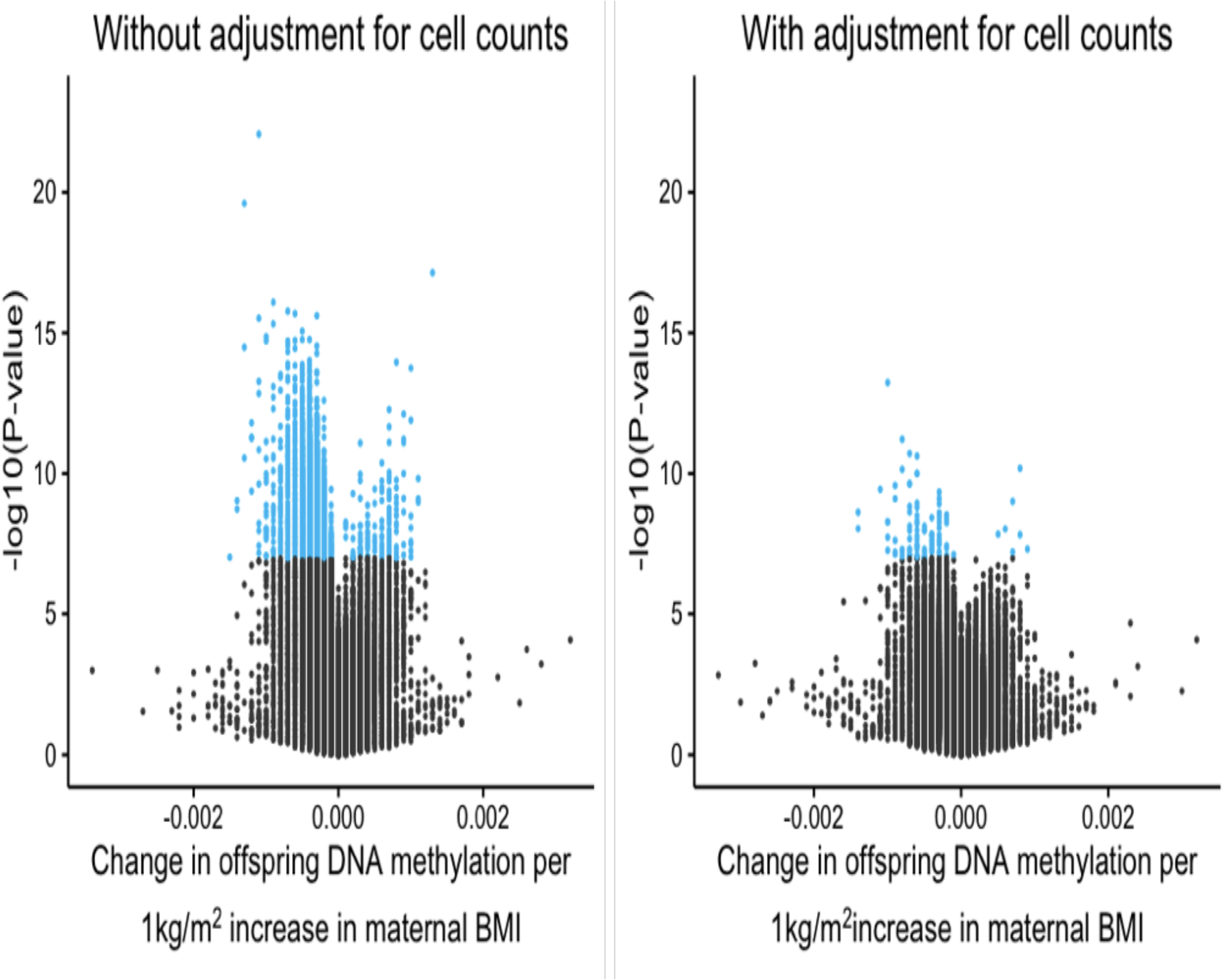
Volcano plots to illustrate the large increase in P-values after adjusting for estimated cell counts. Methylation sites that reached the Bonferroni threshold for multiple testing (1.06* 10-7) are highlighted in blue.

Surprisingly, however, estimated cell proportions were not strongly correlated with maternal BMI in any of the five cohorts that supplied these data (Supplementary File S1 Table S8). Given this, we hypothesised that large changes in estimates might indicate measurement error in estimated cell counts, and that this measurement error might be due to an adult whole blood reference panel being used to estimate cell counts in cord/newborn blood samples. However, we found little evidence for this: Cord blood reference panels by Andrews & Bakulski(28), Gervin et al.(29) and deGoede et al.(30) became available after we had finalised the meta-analysis results. When we used each of these references to estimate cell proportions in ALSPAC cord blood samples, regression coefficients and P-values were similar to those obtained when an adult reference panel was used in this cohort. Of the 86 sites where maternal BMI was associated with newborn methylation before and after adjustment for cell counts in the meta-analysis (P<1.06*10^-7^), 15 were associated with maternal BMI with P<0.05 in ALSPAC when an adult reference panel was used. Of these 15 sites, 12 sites also had P<0.05 when any of the cord blood reference panels were used. The percentage change in estimates between models using the adult and cord blood reference panels was under 10% at 14/15 sites using the Andrews and Bakulski reference (median percentage change in estimates: 4.1), under 10% at 14/15 sites using the Gervin et al. reference (median percentage change in estimates: 3.4) and under 10% at 12/15 using the deGoede reference (median percentage change in estimates: 3.7). Furthermore, cell counts estimated using any of the three cord blood references correlated relatively well with each other (median Spearman’s correlation coefficient: 0.67, range: -0.05 to 0.95), but were not correlated with maternal BMI (median Spearman’s correlation coefficient: 0.007, range: -0.10 to 0.15) (Supplementary File S1 Table S8). Although maternal BMI was not associated with estimated cell proportions in our data, others have observed that maternal BMI is associated with cord blood cellular heterogeneity(31, 32), in addition, some random variability in cell distribution across the range of maternal BMI can be expected. Therefore, we believe that adjustment is appropriate and indeed necessary.

### Further analysis of 86 sites where maternal BMI is associated with newborn DNA methylation both before and after adjustment for cell counts

For further analysis, we selected the 86 sites where maternal BMI at the start of pregnancy was associated with offspring newborn blood DNA methylation both before and after adjustment for estimated cell proportions (Table 3), and performed subsequent analyses using the cell-adjusted model. We used three main strategies to determine the robustness of our findings at these 86 sites:

**Table 3.**
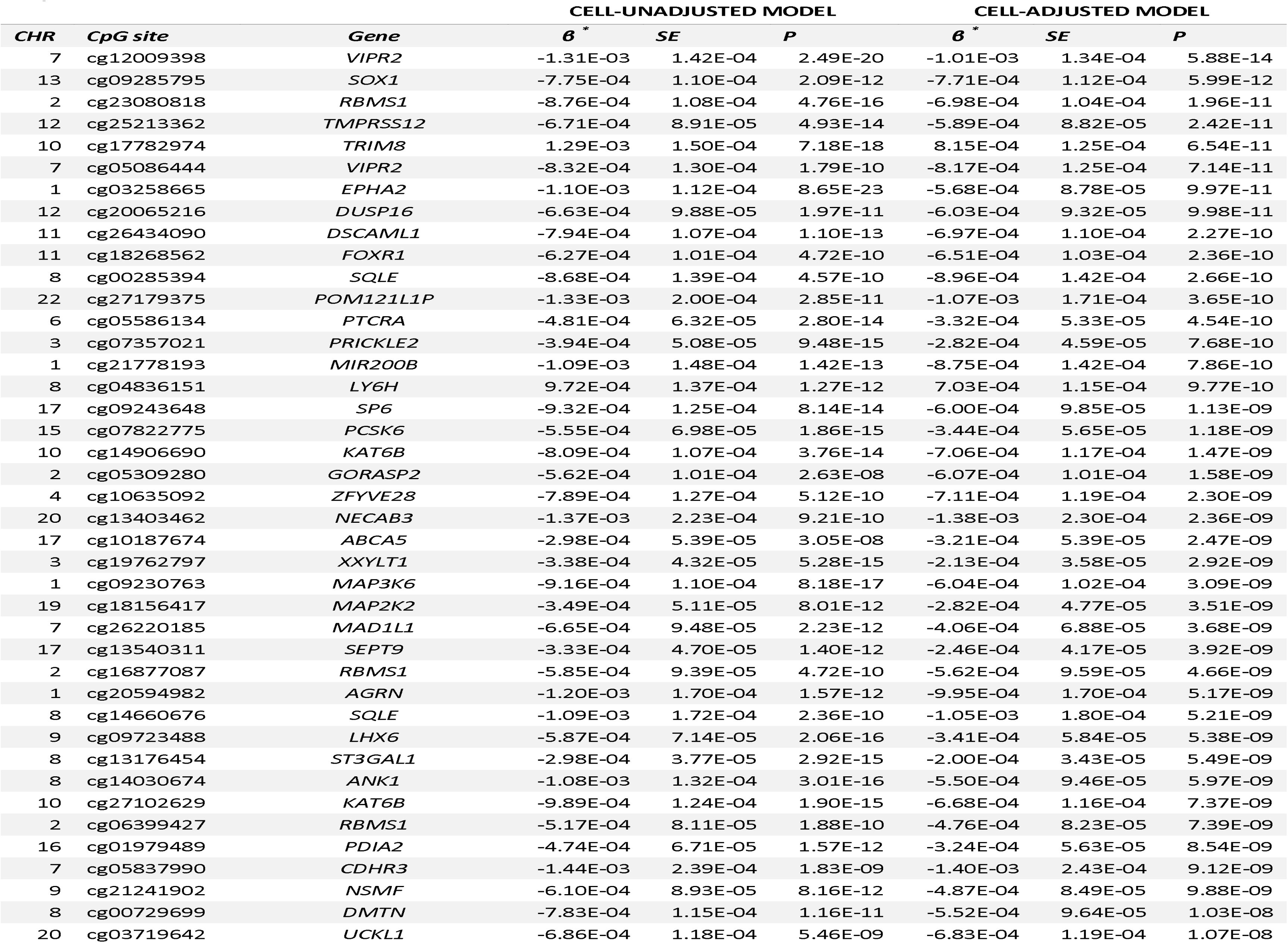
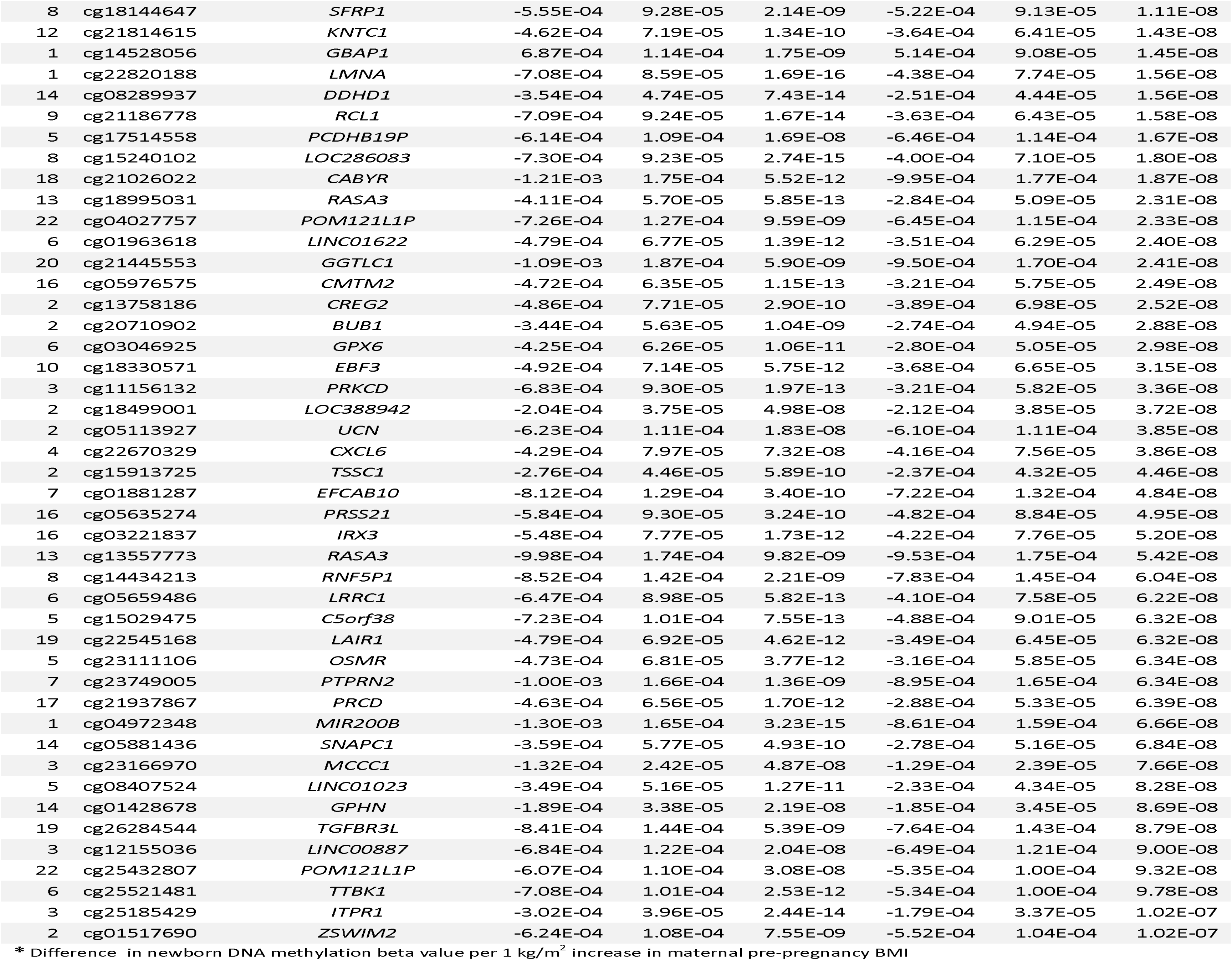
Methylation sites where continuous maternal pre-pregnancy BMI was associated with offspring newborn blood methylation with a Bonferroni-corrected P-value <0.05 (P<1.06*10^-7^) before and after adjustment for cell counts.

Firstly, we assessed inter-study heterogeneity and influence of individual studies. There was weak to moderate heterogeneity at most sites; I^2^ was less than 40% at 57/86 sites (median 31.2%, range 0.0 to 70.6%) and 31/86 sites had a heterogeneity P-value <0.05. In a comparison of estimates from random- and fixed-effects meta-analysis models, the percentage change in estimates was <10% for 72/86 sites (median percentage change in estimates: 2.8). In the random effects model, the largest p-value at the 86 sites was 0.0058 and 20/86 sites had P<1.06*10^-7^, despite lower power compared to the fixed effects model. Forest plots and results of a leave-one-out analysis showed that results from most cohorts agreed on the direction of effect at the 86 top sites and no single cohort consistently had a disproportionately large influence on the meta-analysis (Supplementary File S3).

Secondly, we performed a sensitivity analysis restricting the meta-analysis to 15/19 cohorts comprising participants of European origin only. The results from this sensitivity analysis were consistent with those of the main analysis. The Spearman’s correlation coefficient for regression coefficients was 0.91, and the percentage change in estimates was >10% for 47/86 sites (median percentage change in estimates: 9.7%). While this modest difference could reflect confounding by ancestry, it might also occur because the cohorts of non-European ancestry tended to have a higher mean maternal BMI and were more variable compared to the European ancestry cohorts (Table 1).

Thirdly, we compared the 86 sites to a list of 190,672 probes on the Illumina 450k platform that Naeem et al.(33) suggested might give spurious readings (Supplementary File S1 Table S9). Forty-two sites were on this list: seven located in regions containing SNPs, 11 in regions containing repeat sequences and four in regions where insertions or deletions are found. These sites may be more likely to contain outlier values that influence results, however diptests for multimodality(34) and visual inspection of density plots of methylation beta values in ALSPAC and GOYA did not support this (P>0.05; Supplementary File S3). Additionally, all cohort-specific analyses were conducted using robust linear regression, which is designed to be robust to outliers in the outcome variable (methylation). Other reasons that probes had been flagged by Naeem et al. as potentially problematic were that they hybridise to multiple genomic loci (four sites), did not produce results consistent with those produced by whole-genome bisulfite sequencing (nine sites) and were particularly susceptible to errors in bisulfite conversion (four sites).

### Maternal BMI-associated newborn blood methylation sites are not enriched for certain biological processes or pathways

Maternal BMI-associated newborn blood methylation sites were spread throughout the genome and did not appear to cluster in certain chromosomal regions. The 86 maternal BMI-associated methylation sites are near 77 gene regions, and there were several instances where multiple sites mapped to the same gene: *RBMS1* [3 sites], *POM121L1P* [3 sites], *VIPR2* [2 sites], *SQLE* [2 sites], *RASA3* [2 sites], *MIR200B* [2 sites], *KAT6B* [2 sites]. The list of 77 genes was not enriched for any gene ontology (GO) term (Supplementary File S1 Table S10) or KEGG pathway (Supplementary File S1 Table S11) after FDR-correction for multiple testing, but this analysis was likely underpowered.

### Associations between maternal BMI at the start of pregnancy and newborn DNA methylation were reproduced in the whole blood of adolescents at most sites

In order to assess whether associations at birth are also present in later childhood, four cohorts (BAMSE, IOW birth cohort [IOW F1], PIAMA, and RAINE; total n=1,817 mother-child pairs) contributed results to a meta-analysis of maternal BMI at the start of pregnancy and methylation in the whole blood of adolescent offspring (age range: 15 to 18 years, weighted mean: 17 years). Cohorts are summarised in Table 4. These cohorts were completely independent of those that contributed results to the newborn analysis, therefore we were able to assess reproducibility of our newborn results later in life. All models discussed here were corrected for estimated cell counts. Full results are provided in Supplementary File S1 Table S12.

**Table 4.** Characteristics of each cohort included in the meta-analysis of the effect of maternal pre-pregnancy BMI on off spring DNA methylation at adolescence

| COHORT | N | MEAN MATERNAL BMI (SD) | MEAN MATERNAL AGE (SD) | MEAN<br>ADOLESCENT AGE<br>(SD) | ETHNICITY |
| --- | --- | --- | --- | --- | --- |
| BAMSE | 221 | 23.2 (3.4) | 31.2 (4.3) | 16.6 (0.3) | European |
| IOW F1 | 279 | 24.4 (4.0) | 27.3 (5.2) | 18.0 (0.0) | European |
| PIAMA | 583 | 22.6 (3.1) | 30.9 (3.7) | 16.3 (0.2) | European |
| RAINE | 734 | 22.4 (4.4) | 29.1 (5.8) | 17.3 (0.6) | European |
| **Meta-analysis** | **1817** |  |  |  |  |

There was evidence for reproducible associations at most of the 86 sites: the direction of association at adolescence was the same as that at birth for 72/86 sites (Spearman correlation coefficient: 0.67). Twenty-two of these 72 sites had a P-value <0.05 at adolescence, despite the much smaller sample size. Although no associations survived correction for multiple testing at 86 sites, 22/72 sites with nominal P-values <0.05 is higher than the 5% expected by chance alone (Kolmogorov P=3.3 *10^-16^). Across the 72 sites where effects were in the same direction, the effect estimates in the adolescence analysis were a median of 2.25 times smaller (i.e. closer to the null) than the effect estimates in the newborn analysis (range: 2889 times smaller to 1.35 times larger) but at some sites, estimates at both time points were remarkably similar (Figure 4). It is also of particular note that six of the top ten sites with the largest effect size were the same at birth and adolescence. These sites were cg05837990 (*CDHR3*), cg13403462 (*ACTL10/NECAB3*), cg27179375 (*POM121L1P*), cg12009398 (*VIPR2*), cg20594982 (*AGRN*) and cg21445553 (*GGTLC1*). One of the top ten sites with the smallest P-values was also common to both analyses: cg05086444 (*VIPR2*).

**Figure 4.**
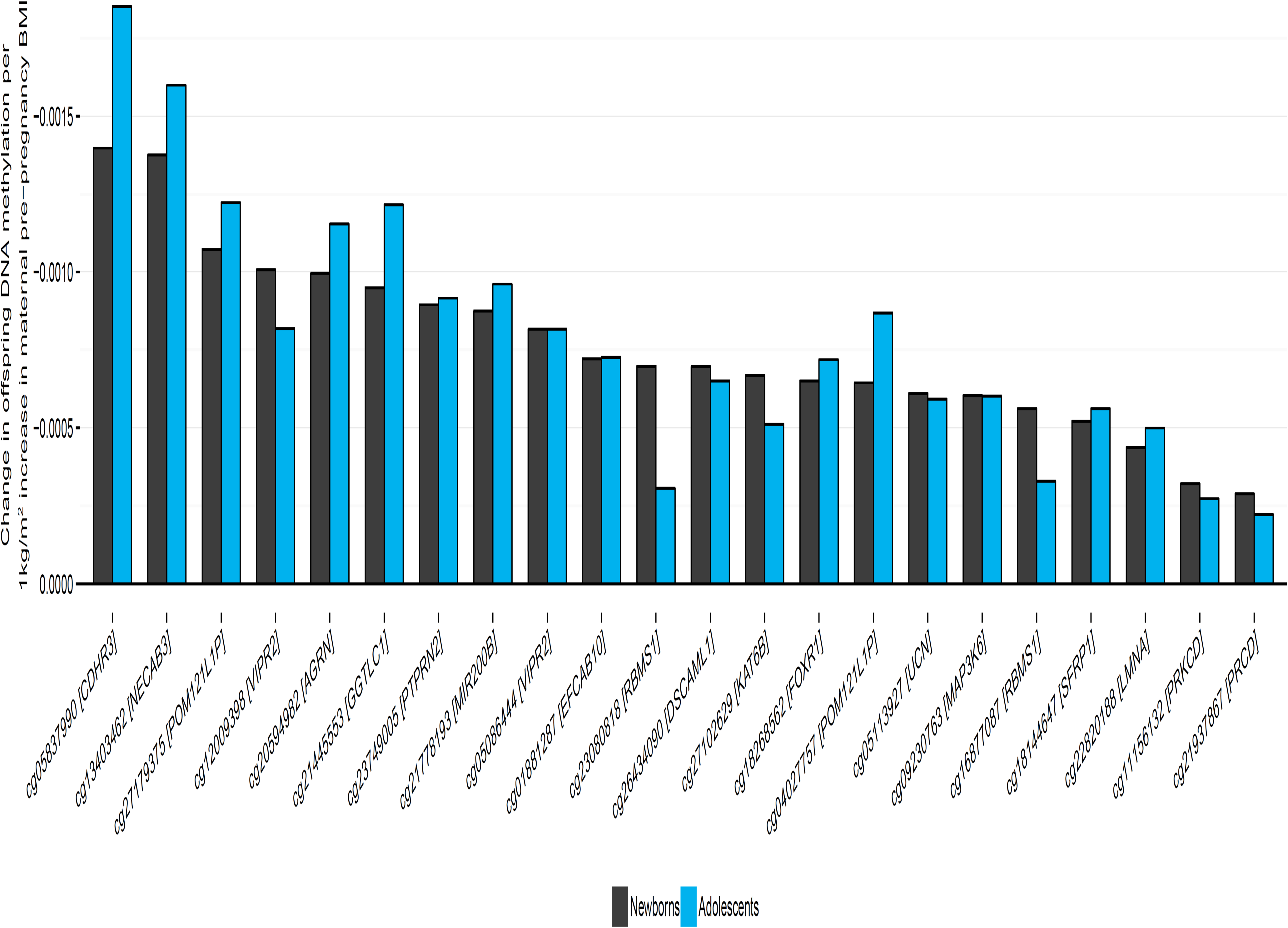
Comparison of estimates of the effect of maternal BMI on offspring DNA methylation at birth and at adolescence. Of the 86 sites where maternal BMI at the start of pregnancy was associated with newborn blood methylation, 72 had the same direction of association in the analysis of adolescents. Plotted here are the 22/86 methylation sites with a P-value<0.05 in the analysis of adolescents, ordered by effect size in newborns.

### Negative control design supports a causal intrauterine effect of maternal BMI on newborn blood methylation at nine sites

We used a negative control design(7) in an attempt to disentangle a potential causal, intrauterine effect of maternal BMI on newborn blood methylation from the effect of confounding by shared genetics or postnatal environment. The logic is that paternal and maternal exposures may both be associated with offspring methylation due to shared familial confounding factors or by inheritance of parental genotypes, but paternal BMI would not normally be expected to affect the intrauterine environment. Therefore, if there is a causal intrauterine influence, only maternal BMI would be expected to be independently associated with methylation. Evidence for an intrauterine effect is stronger where estimates for associations between maternal BMI and offspring DNA methylation are greater than the equivalent estimates for paternal BMI., whereas consistent maternal and paternal estimates provides evidence for confounding by genetic or shared postnatal environmental factors.

It is also important to adjust the maternal estimate for paternal BMI, and vice versa, because maternal and paternal BMI are somewhat correlated due to assortative mating. For example, in the cohorts that contributed to this study, Spearman’s correlation coefficients between maternal and paternal BMI ranged from 0.18 to 0.25 (p<0.001).

Seven cohorts contributed results to this negative control analysis: ALSPAC (n=619), CHAMACOS (n=180), Generation R (n=829), GOYA (n=422), MEDALL (INMA and EDEN pooled n=316), NHBCS (n=96) and RICHS (n=92). The total number of families included in the meta-analysis of the mutually adjusted models was 2,554. Results for all models are provided in Supplementary File S1 Table S13.

Based on the above criteria, we found some evidence for a causal intrauterine effect of maternal BMI on newborn blood methylation at some sites: At 64 of 86 sites, the paternal and maternal effect estimates were in the same direction, i.e. we could be more certain that no independent paternal-specific effect exists. At 40 of these 64 sites, the maternal BMI estimate was greater than the paternal BMI estimate after mutual adjustment (median 2.19 times greater, range 1.01 to 142.4 times greater). At nine of these 40 sites, there was some evidence of heterogeneity between the mutually adjusted maternal and paternal BMI estimates (I^2^>40; Table S14). These criteria were used to define support for a possible maternal specific, intrauterine effect. Therefore, at 77/86 sites, evidence from this negative control study was more supportive of the association between maternal BMI and newborn blood methylation being explained by genetic or shared prenatal environmental factors than a causal intrauterine effect. Figure 5 displays the results for the 20 sites where the mutually adjusted maternal and paternal BMI estimates were in the same direction, with the maternal effect being larger than the paternal effect and having a P-value <0.05 (Figure 5).

**Figure 5.**
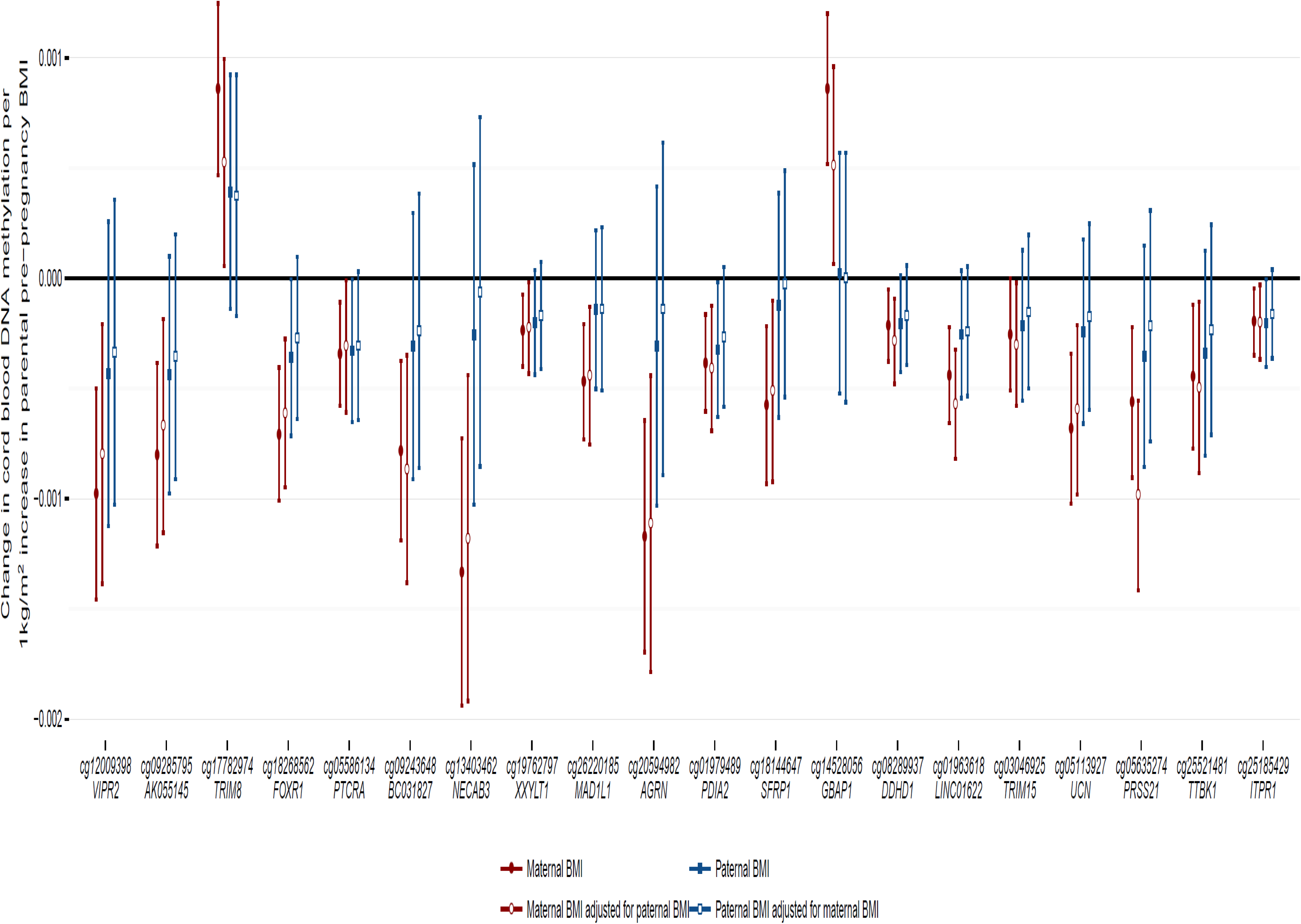
Comparison of estimates of the effect of maternal and paternal BMI on newborn DNA methylation. Of the 86 sites where maternal BMI at the start of pregnancy was associated with newborn blood methylation, we found 20 sites (plotted here) where the estimated effect of maternal BMI, adjusted for paternal BMI, had a P-value<0.05 and was in the same direction and greater than the estimated effect of paternal BMI, adjusted for maternal BMI. Sites are ordered by P-value in the full maternal BMI meta-analysis.

### meQTLs at maternal BMI-associated cord blood methylation sites provide further support for confounding by genetics at four sites

To explore the genetic influence on DNA methylation at the 86 maternal BMI-associated cord blood methylation sites, we performed a look-up in an online catalogue of methylation quantitative trait loci (meQTL) that were previously identified using ALSPAC data(35). We identified 821 meQTLs where genetic variation was associated with cord blood DNA methylation at 27/86 sites with P<1*10^-7^. Of these 821 meQTLs, 68 were within 1Mb of the methylation site (cis) and 753 were outside of this window (trans).

If an meQTL is also associated with maternal BMI, this could suggest that the association between maternal BMI and newborn methylation is confounded by shared genetics. Of the 821 identified meQTLs, data for 225 were available in the results of the largest adult BMI GWAS meta-analysis to date, conducted by the GIANT consortium (36). Of these, 17/225 were nominally associated (P<0.05) with BMI in GIANT. These 17 meQTLs were associated with cis methylation at four CpGs: 11 with cg03258665 *(EPHA2*), four with cg00285394 (*SQLE*), one with cg03719642 (*UCKL1*) and one with cg18268562 (*FOXR1*). Therefore, there is some evidence that associations between maternal BMI and methylation at these four are confounded by shared genetics. For most of the meQTLs, the associations SNP-BMI and SNP-methylation were in opposite directions. Thus, the same effect allele was associated with higher BMI (effect estimates ranging 0.007 to 0.015) and lower methylation (effect estimates ranging -0.523 to -0.235). Only in the rs8567-cg03719642 association was the effect allele associated with lower BMI (effect estimate: -0.012) and higher methylation (effect estimate: 0.287).

### Using a combination of evidence, we identified eight sites where maternal BMI may have a causal intrauterine effect on newborn blood methylation

As described above, by employing a negative control design, we found nine sites where the estimated effect of maternal BMI was stronger than that of paternal BMI. One of these sites (cg18268562 at *FOXR1*) is an meQTL that was nominally associated with BMI in GIANT. Therefore, we find strongest support for a causal intrauterine effect of maternal BMI at the start of pregnancy on newborn blood methylation at just eight sites (Table 5). At the remaining 78 of our top 86 sites, the apparent associations between maternal BMI and newborn blood methylation might be more appropriately explained by shared mother-offspring genetic and postnatal environmental factors. These findings are summarised in Supplementary File S1 Table S14.

**Table 5.**
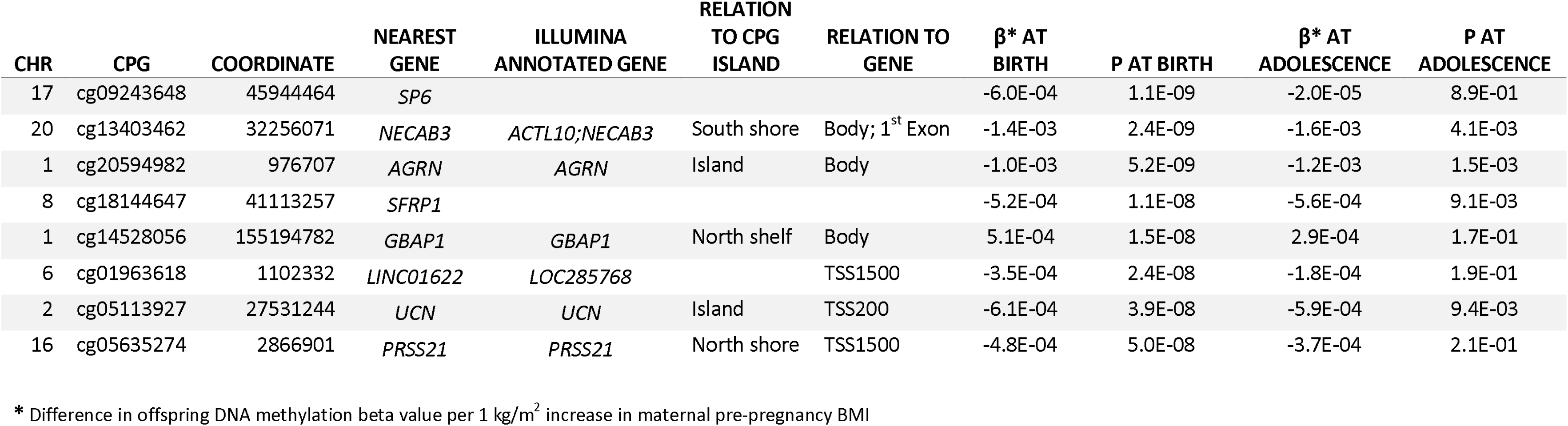
A summary of the 8 sites where there is strongest evidence for a causal intrauterine effect of maternal BMI on newborn blood DNA methylation

| CHR | CPG | COORDINATE | NEAREST GENE | ILLUMINA ANNOTATED GENE | RELATION TO CPG ISLAND | RELATION TO GENE | $\beta^*$ AT BIRTH | P AT BIRTH | $\beta^*$ AT ADOLESCENCE | P AT ADOLESCENCE |
| --- | --- | --- | --- | --- | --- | --- | --- | --- | --- | --- |
| 17 | cg09243648 | 45944464 | *SP6* |  |  |  | -6.0E-04 | 1.1E-09 | -2.0E-05 | 8.9E-01 |
| 20 | cg13403462 | 32256071 | *NECAB3* | *ACTL10;NECAB3* | South shore | Body; 1<sup>st</sup> Exon | -1.4E-03 | 2.4E-09 | -1.6E-03 | 4.1E-03 |
| 1 | cg20594982 | 976707 | *AGRN* | *AGRN* | Island | Body | -1.0E-03 | 5.2E-09 | -1.2E-03 | 1.5E-03 |
| 8 | cg18144647 | 41113257 | *SFRP1* |  |  |  | -5.2E-04 | 1.1E-08 | -5.6E-04 | 9.1E-03 |
| 1 | cg14528056 | 155194782 | *GBAP1* | *GBAP1* | North shelf | Body | 5.1E-04 | 1.5E-08 | 2.9E-04 | 1.7E-01 |
| 6 | cg01963618 | 1102332 | *LINC01622* | *LOC285768* |  | TSS1500 | -3.5E-04 | 2.4E-08 | -1.8E-04 | 1.9E-01 |
| 2 | cg05113927 | 27531244 | *UCN* | *UCN* | Island | TSS200 | -6.1E-04 | 3.9E-08 | -5.9E-04 | 9.4E-03 |
| 16 | cg05635274 | 2866901 | *PRSS21* | *PRSS21* | North shore | TSS1500 | -4.8E-04 | 5.0E-08 | -3.7E-04 | 2.1E-01 |

\* Difference in offspring DNA methylation beta value per 1 kg/m<sup>2</sup> increase in maternal pre-pregnancy BMI

## Discussion

We found that maternal BMI at the start of pregnancy is associated with small variation in newborn blood DNA methylation at 86 sites throughout the genome, after adjusting for cell proportions. At around a quarter of these 86 sites, we found nominal associations between maternal pre-pregnancy BMI and DNA methylation in an independent cohort of adolescents, sometimes with remarkably consistent effect sizes to those found in neonates. However, when we employed two causal inference strategies, we found supporting evidence for a causal intrauterine effect at only eight sites. Taken together, our results suggest that the effects of maternal pre-pregnancy adiposity on neonatal blood DNA methylation are primarily related to variations in the cellular distributions in cord blood, as well as shared environment and genetic variation. Although there may be a causal intrauterine effect at some sites, the biological significance of such small effects is unclear.

Our findings are in contrast to some previous studies that have reported strong associations between maternal BMI/adiposity and DNA methylation in neonates(21, 24–26, 37, 38). However, in these smaller studies there has been a lack of consistency in terms of the specific loci identified. Although we replicated, at look-up level of significance, an inverse association between maternal BMI and newborn blood methylation at cg01422136 (*ZCCHC10*) that was reported in a study of African American and Haitian mother-child pairs from the Boston Birth Cohort(24), this association was not epigenome-wide significant in our study (P=0.0016). We did not replicate specific associations reported in other previous studies of maternal BMI and newborn blood methylation, including some that were reported in individual studies from the PACE consortium(21, 25, 26, 37, 38). This lack of consistency highlights the potential presence of false positive findings in small EWAS studies and the importance of meta-analysis for improving power and reproducibility.

The 86 Bonferroni-significant sites were robust across cohorts and after adjustment for cell proportions, so they are unlikely to have arisen due to chance, study-specific biases or technical aspects of the array, which should be independent of our exposure. However, effect sizes were very small; all were less than a 0.15% change in methylation per one-unit increase in maternal BMI. The biological significance of such small effects is unclear and could not be further explored in this study due to lack of genome-wide data on downstream gene and protein expression. One reason we may not have observed larger effect sizes is that the studied cohorts consisted mostly of women whose weight fell within the WHO BMI category of normal weight. Perhaps the largest effects only exist at the extremities of the BMI distribution, as is the case with some other maternal BMI-associated offspring phenotypes, including offspring BMI(6). However, we also found relatively small effects in our binary exposure model comparing methylation in offspring of women who were overweight or obese to methylation in offspring of women who were normal weight at the start of pregnancy.

Without integration with gene expression data, it is impossible for us to truly infer (either way) whether maternal BMI-associated variation in methylation at our 86 sites is functionally important. The 77 mapped genes were not enriched for any GO term or KEGG pathway, which could suggest that there is little or no significant biological effect. However, this analysis was likely underpowered and it is worth noting that, individually, some of the 77 genes that map to our 86 sites have functions that could potentially link maternal adiposity to offspring health outcomes, either through shared genetic factors or an epigenetic effect on gene regulation. These may be useful candidates for future studies that are better placed to explore the biological significance of the methylation sites we have identified. For example, GWAS studies have identified that variants at some of our differentially methylated loci are associated with adiposity-related traits: total energy total energy expenditure (*CDHR3*(41)), energy intake (*PTPRN2*(41)), lipoprotein-a levels (*DSCAML1*(42)), adiponectin levels (*CREG2*(43)), and type 2 diabetes (*ANK1, RBMS1*(44–46)). In studies of DNA methylation, greater whole blood methylation at cg17782974 (*TRIM8*) was associated with higher BMI in elderly participants in the Lothian Birth Cohort study(47) and higher maternal BMI in our study. Another 450k study found that several sites at *PTPRN2* were hypermethylated in subcutaneous adipose tissue of women before gastric-bypass compared to the same women after gastric-bypass and associated weight-loss(48), whereas we found that higher maternal BMI at the start of pregnancy was associated with hypomethylation at *PTPRN2* in newborn blood. We also found that higher maternal BMI at the start of pregnancy was associated with lower newborn methylation at a site (cg03221837) near *IRX3.* More copies of the risk allele at the obesity-associated SNP *FTO* is associated with higher blood expression of *IRX3* in humans, and *IRX3*-deficient mice have been shown to have a 25-30% reduction in body weight(49). However, it is important to note that although *IRX3* was the nearest gene to the maternal BMI-associated methylation site in our study, the site was actually 299,591 bp downstream from the gene. Finally, we were particularly interested to find two sites (cg12009398, cg05086444) on the gene body of *VIPR2* where greater maternal BMI was associated with lower methylation. The associations were consistent in adolescents, with P-values <0.008, although we did not find any evidence that the associations were causal. *VIPR2* encodes vasoactive intestinal peptide receptor 2 (VIPR2), which functions as a neurotransmitter and as a neuroendocrine hormone. A GWA analysis in 1,000 participants found that the vasoactive intestinal peptide (VIP) pathway was strongly associated with fat mass and with BMI, suggesting that the VIP pathway may play an important role in the development of obesity(52). In a study using the 450k array, lower *VIPR2* methylation was found in the saliva of children with attention deficit hyperactivity disorder (ADHD), relative to controls(53), albeit at different sites than those identified in the present study. Given previously identified associations between maternal BMI and offspring ADHD(54–57), further work is warranted to explore the extent to which *VIPR2* gene function (driven either by genetic variation or regulation by methylation) might explain associations between maternal adiposity and neurodevelopment of the offspring.

Of the 86 sites where maternal BMI was associated with methylation in the blood of newborns, 72 showed the same direction of association in the blood of an independent smaller sample of adolescents. At some sites, effect estimates were remarkably consistent between the two age groups. Of particular note, six of the top 10 sites with the largest effect size in the cell-adjusted newborn analysis also had the largest effect size amongst adolescents. This consistency from birth to adolescence could be explained as either i) an intrauterine influence of maternal pre-pregnancy BMI on variation in offspring DNA methylation that persists to adolescence, ii) confounding by shared familial genetic and/or environmental influences on maternal BMI and offspring methylation that remain stable over time, or iii) the possibility that both maternal pre-/early-pregnancy and the adolescent’s own BMI have independent effects on the child’s methylation. We did not adjust for adolescent’s BMI because that may introduce a collider that would bias the association between shared familial factors and maternal BMI away from the null.

We were interested in whether the 86 maternal BMI-associated sites represented a causal intrauterine effect of maternal adiposity on offspring methylation, or if associations were better explained by confounding by shared environment or genetics. By employing a negative control design, we found nine sites where the estimated effect of maternal BMI was larger than that of paternal BMI, after mutual adjustment. Maternal and paternal BMI were not strongly correlated in any of the cohorts that took part in this analysis (Spearman’s R ranging 0.13 to 0.25), so collinearity in the mutually adjusted models is unlikely to bias interpretation of results. This is supported by the observation that standard errors did not increase substantially between the unadjusted and adjusted models. At one of the nine sites (cg18269562 mapping to *FOXR1*), cord blood methylation has previously been strongly associated (P<1*10^-7^) with common genetic variants(35). This meQTL was also nominally associated (P<0.05) with BMI according to the GIANT consortium adult BMI GWAS metaanalysis(36, 58). We considered that the association between maternal BMI and newborn methylation at this site was likely driven by a shared genetic effect. Therefore, we could be most confident of a causal intrauterine effect of maternal adiposity on methylation of blood DNA in newborns at only 8/86 sites. At the remaining 78/86 sites, shared genetic and/or prenatal environmental factors, which would be expected to be the same whether the exposure were maternal or paternal BMI, may have larger influences on newborn blood methylation than maternal BMI at the start of pregnancy.

Our findings are in line with studies reporting that a large proportion of variation in DNA methylation is explained by genetics. One study estimated that at around 50% of CpG sites on the Illumina 450k array methylation has a substantial genetic component(59). Another study of DNA methylation using the same platform in 237 neonates found that, of 1,423 genomic regions that were highly variable across individuals, 25% were best explained by genotype alone and 75% by an interaction of genotype with different *in utero* environmental factors (including maternal BMI)(60). These studies, along with our own, highlight complex relationships between genetic inheritance, intrauterine environmental exposures and offspring epigenetics. In light of this, we recommend that where the exposure is genetically heritable, extra care should be taken to avoid over-interpreting EWAS results as representing causal environmental effects(61). Causal analysis techniques, such as the negative control and meQTL analyses conducted in this study, will be useful in this regard.

Regardless of whether maternal BMI has a biologically significant, causal effect on newborn blood DNA methylation, the robust, and seemingly persistent, associations we identified in our study suggests that, as has been shown for maternal smoking(62), blood DNA methylation could be a useful indicator of maternal BMI during pregnancy. Such an indicator would be useful in studies where maternal BMI data is missing. Likewise, newborn blood methylation at maternal BMI-associated sites might also be predictive of offspring outcomes, capturing both genetic and environmental influences of maternal adiposity.

Although our findings suggest no strong effect of maternal pre-pregnancy adiposity (as measured by BMI) on offspring methylation in blood, this does not preclude the possibility that there is an effect of maternal adiposity measured in different ways and/or on offspring methylation in different tissues. It will be interesting to explore in further work how maternal adiposity-associated exposures *during* pregnancy, such as gestational weight gain, maternal hypertension and hyperglycemia, influence offspring DNA methylation. Such pregnancy exposures may be more likely to have a pronounced intrauterine effect on offspring methylation and/or developmental programming of health outcomes than maternal adiposity at the start of pregnancy. Although previous studies in ALSPAC(21) and MoBa(63) did not identify any sites where gestational weight gain was associated with cord blood methylation, the question should be revisited in a consortium context. Further exploration is also warranted to assess the degree to which methylation in blood correlates with that in other tissues. DNA methylation shows strong tissue-specificity, for example, one study found that BMI was associated with DNA methylation in adipose tissue, but not in peripheral blood leukocytes(64). Conversely, a large EWAS found that BMI was associated with methylation at *HIF3A* in both blood and adipose tissues(65). The causal effect of maternal BMI on newborn methylation may be stronger in tissues other than blood. However, we note that in the context of this study, offspring blood might be considered a mechanistically relevant tissue: blood cellular heterogeneity and leukocyte methylation are strongly associated with inflammation, which is considered chronic amongst those with obesity.

There are several strengths to our study, including the large sample size comprised of established cohorts, the use of robust statistical methods, the comprehensive analysis of results and the application of causal inference techniques. Potential limitations include: i) adiposity is a complex trait that is only crudely and indirectly measured by BMI, therefore an investigation of more specific measures of adiposity might yield different results, ii) cohorts collected data on BMI in different ways (measured/self-reported) at different times (prepregnancy/early pregnancy). However, measured and self-reported BMI before and during early pregnancy are strongly correlated(66), so we do not believe this will bias our results substantially. iii) The analysis was completed before the widespread availability of any cord blood reference panels for estimations of cell counts, so all cohorts used an adult whole blood reference panel, which may introduce measurement error in cell count estimates(28). However, in ALSPAC, one of the largest participating cohorts, we found that adjusting for cell counts generated using any one of three recently released cord blood reference panels produced results consistent with those produced using the adult whole blood reference. Nevertheless, we consider that there is likely to be at least some degree of residual influence of cell heterogeneity in our results. iv) We had very limited data with repeat measures in the same individuals at birth and adolescence, so we did not explore change in methylation over time in a longitudinal model. v) Cohorts used different methods to normalise data. However, a previous PACE analysis(67) found that results obtained using raw betas were similar to those obtained using normalized betas generated with various methods, which indicates that this did not impact the inferences drawn from the meta-analysis, and at any rate, bias would tend to limit power rather than introduce spurious <u>associations. vi</u>) Although we have presented two lines of evidence (consistent maternal and paternal estimates and the presence of meQTLs) that provide support for a genetic component in explaining associations between maternal BMI and newborn blood methylation at some sites, we were unable to formally quantify the relative contribution of genetics and the intrauterine environment. Techniques that attempt to do so, such as M-GCTA(68), require genetic and methylation data on larger sample sizes than were available in any individual cohort. vii) The Illumina 450k array only covers 1.7% of CpG sites on the human genome, and most of these are located in promoter regions. We found robust associations between maternal BMI and newborn DNA methylation despite this low coverage and bias. We therefore encourage more studies on this topic using more advanced EWAS platforms (such as the Illumina EPIC array). viii) Finally, it is possible that epigenetic markers other than DNA methylation in cord blood may be more closely associated with maternal BMI at the start of pregnancy, but this was not explored in this study.

In conclusion, in this well-powered study, we observed robust associations between maternal pre/early-pregnancy BMI and DNA methylation at 86 sites in the blood of newborns, some of which were reproduced in adolescents. However, effect sizes were very small, there was no evidence of biological functional enrichment, and causal inference strategies provided support for causal effects at just 8/86 sites. This study highlights that although some small studies report strong associations between prenatal exposures and epigenetics, large-scale collaborative efforts are necessary to identify robust associations, and causal inference strategies are needed to assess whether such associations are likely to be explained by a direct intrauterine effect or more likely due to genetic or shared environmental factors.

## Methods

Figure 1 gives an outline of the design of this study.

### Participating cohorts

A total of 23 independent cohorts participated. Detailed methods for each cohort are provided in the Supplementary Material (Supplementary File S2) and summarised in Supplementary File S1 Table S1.

Nineteen cohorts participated in the meta-analysis of maternal BMI at the start of pregnancy and newborn blood DNA methylation: The Avon Longitudinal Study of Parents and Children (ALSPAC)(69–71); two independent datasets from the Californian Birth Cohort (CBC_Hispanics and CBC_Caucasians) (72); Center for the Health Assessment of Mothers and Children of Salinas (CHAMACOS); Early Autism Risk Longitudinal Investigation (EARLI) (73); the Genome-Wide Population-based Association Study of Extremely Overweight Young Adults (GOYA), which is a sample from the Danish National Birth Cohort (74, 75); Groningen Expert Center for Kids with Obesity (GECKO); Generation R (GENR)(76); Genetics of Glycemic Regulation in Gestation and Growth (GEN3G)(77); the Isle of Wight Birth Cohort third generation (IOW F2) (78); two cohorts from the FP7 project Mechanisms of the Development of Allergy (MEDALL), INfancia y Medio Ambiente (INMA) (79) and a study on the pre- and early postnatal; determinants of child health and development (EDEN)(80), were pooled and analysed as a single cohort referred to as MEDALL; three independent datasets from the Norwegian Mother and Child Cohort Study (MOBA1, MOBA2, MOBA3)(81, 82); the Norway Facial Clefts Study (NFCS), the Newborn Epigenetic Study (NEST)(83, 84); the New Hampshire Birth Cohort Study (NHBCS); the Rhode Island Child Health Study (RICHS)(85) and Project Viva (Viva).

An additional four independent cohorts participated in the meta-analysis of maternal BMI at the start of pregnancy and offspring whole blood DNA methylation at adolescence (ages 15-18): the Children Allergy Milieu Stockholm Epidemiology cohort (BAMSE) (86), IOW birth cohort second generation (IOW F1), the Prevention and Incidence of Asthma and Mite Allergy birth cohort (PIAMA), the Western Australia Pregnancy Cohort (RAINE).

All cohorts acquired ethics approval and informed consent from participants prior to data collection through local ethics committees. Full details are provided in Supplementary File S2.

### Maternal BMI at the start of pregnancy

In each cohort, maternal BMI (weight (kg)/height (m2)) was calculated from either self-reported or measured height and weight, either before pregnancy or early in the first trimester (Supplementary File S1 Table S1). Cohorts were asked to double check values ≥5 standard deviations from the mean to ensure that they were not data entry errors. Primarily, we were interested in the effects of maternal BMI as a continuous variable, but also investigated World Health Organization categories of maternal overweight or obesity (≥25.0 kg/m2), and underweight (<18.5 kg/m2), compared to a normal weight reference group (18.5-24.9 kg/m2).

### Covariates

All cohorts ran models adjusted for maternal age (years), maternal social class (variable defined by each individual cohort), maternal smoking status (the preferred categorization was into three groups: no smoking in pregnancy, stopped smoking in early pregnancy, smoking throughout pregnancy, but a binary categorization of any versus no smoking was also acceptable) and parity (the preferred categorization was into two groups: no previous children, one or more previous children). We did not adjust for or stratify by sex of the child because sex cannot be a true confounder of any association between maternal pre-pregnancy BMI and offspring methylation; although it has a large influence on methylation, it cannot feasibly alter pre-pregnancy BMI. Furthermore, because the intrauterine hormonal environment is likely to be different for males and females, and could also be influenced by maternal BMI, we would risk introducing collider bias by adjusting for sex, which would be strongly correlated with sex-associated hormonal environment on the causal pathway between maternal BMI and methylation.

Each cohort also adjusted for technical covariates using methods suitable for that cohort (Supplementary File S1 Table S1). Certain cohorts also included additional covariates to correct for study design/sampling factors where needed (Supplementary File S1 Table S1). For GOYA, which is a case-control study where case mothers have a BMI>32kg/m^2^ and control mothers have a BMI anywhere within the normal distribution, we restricted the continuous maternal BMI models to a randomly selected sub-group with a normal BMI distribution to avoid confounding by substructure. Binary comparison models were run using the whole GOYA cohort with no additional adjustment for substructure.

We hypothesised that BMI might influence newborn blood cellular composition, so each cohort additionally adjusted for cell proportions by including the estimated variables as covariates. All cohorts independently estimated cell counts using the *estimateCellCounts* function in the *minfi* R package, which is based on the method developed by Houseman(87, 88). The cohort-specific analyses, as well as the meta-analyses, were completed before a cord blood reference set was widely available, so cohorts used an adult whole blood reference to estimate cell counts(89). This estimated the proportion of B-cells, CD8+ T-cells, CD4+ T-cells, granulocytes, NK-cells and monocytes in each sample. NHBCS, RICHS and Project Viva included five estimated cell types (omitting granulocytes) and all other cohorts included six. When cord blood references became available(28, 30, 90), a sensitivity analysis was run in ALSPAC adjusting for cell proportions estimated using these reference sets. One of these reference sets includes nucleated red blood cells, which can contribute greatly to cord blood DNA methylation profiles(28).

### Methylation measurements and quality control

Each cohort conducted its own laboratory measurements. DNA from newborn or adolescent blood samples underwent bisulfite conversion using the EZ-96 DNA Methylation kit (Zymo Research Corporation, Irvine, USA). For all cohorts, DNA methylation was measured using the Illumina Infinium® HumanMethylation450 BeadChip assay(91, 92) at Illumina or in cohort-specific laboratories. Each cohort also conducted its own quality control and normalisation of methylation data, as detailed in the Supplementary Methods (Supplementary File S2) and summarised in Supplementary File S1 Table S1. In all analyses, cohorts used normalised, untransformed beta-values, which are on a scale of 0 (completely unmethylated) to 1 (completely methylated).

### Cohort-specific statistical analyses

Each cohort performed independent epigenome-wide association studies (EWAS) according to a common, pre-specified analysis plan. Models were run using M-type multiple robust linear regression (rlm in the MASS R package(93)) in an attempt to control for potential heteroscedasticity and/or influential outliers in the methylation data. In the primary analysis, continuous maternal BMI at the start of pregnancy was modelled as the exposure and offspring individual CpG-level methylation (untransformed beta-values) was modelled as the outcome, with adjustment for covariates and estimated cell counts. In secondary models, we modelled the exposure as binary variables comparing WHO BMI categories to a normal weight reference group. We also explored the impact of cellular composition by comparing models run with and without adjustment for estimated cell counts.

### Meta-analysis

Cohorts uploaded their EWAS results files to a server at the University of Bristol, where we performed fixed-effects meta-analysis weighted by the inverse of the variance with METAL(94). A shadow meta-analyses was also conducted independently by authors at the Erasmus University in Rotterdam to minimise the likelihood of human error. All downstream analyses were conducted using R version 2.5.1 or later(95). We excluded control probes (N=65), and probes mapped to the X (N=11,232) or Y (N=416) chromosomes. This left a total of 473,864 CpGs measured in at least one cohort (218,350 (46%) of these were measured in all 19 cohorts, 393,986 (83%) were measured in at least 18 cohorts). Multiple testing was accounted for using the Bonferroni method. CpGs with a Bonferroni-corrected P-value <0.05, i.e. P<1.06*10^-7^, in both the cell proportion-unadjusted and cell proportion-adjusted models were taken forward for further analysis. To assess heterogeneity, we generated forest plots, and ran random effects models and “leave-one-out” analyses using the metafor R package(96). We compared our Bonferroni-siginificant probes to a list of potentially problematic probes published by Naeem et al. We did not remove these probes as this would risk removing potentially interesting effects. However, we tested whether these probes contained large numbers of outlying values by performing dip tests s(97) for multimodality using the diptest package(34), where a p>0.05 suggests the distribution is unimodal. Kolmogorov-Smirnov tests were used to compare the distribution of P-values to that expected by chance and were conducted using the core R function ks.test().

### Enrichment and functional analysis

Sites were annotated using the IlluminaHumanMethylation450k.db R package(98), with enhanced annotation for nearest genes within 10Mb of each site, as previously described(67). These annotations were then updated using the R package mygene(99). Gene Ontology (GO) and Kyoto Encyclopedia of Genes and Genomes (KEGG) enrichment analyses were performed using the missMethyl R package(100). This takes into account the differing number of sites associated with each gene on the 450k array. P-values for enrichment were adjusted for multiple testing using the FDR method.

### Reproduction of maternal BMI-related differential DNA methylation in adolescence

Four cohorts independently performed robust linear regression to assess the association between maternal BMI at the start of pregnancy and (mixed gender) adolescent whole blood DNA methylation. Each of these cohorts ran models adjusted for maternal smoking, maternal age, socioeconomic status, parity during the index pregnancy and estimated cell counts. Results were uploaded to the server at the University of Bristol where they were summarised using fixed effects meta-analysis in the metafor package(96). A look-up of maternal BMI-related sites identified in the newborn meta-analysis (n=86 with P<1.06*10^-7^ in the cell-adjusted and cell-unadjusted models) was performed and FDR correction applied to account for multiple testing. These “reproduction” cohorts were completely independent from the original “discovery” cohorts.

### Negative control design

In an attempt to examine a potential causal effect of maternal BMI on newborn blood methylation at identified sites, we used a negative control design(7). In this analysis, estimates for associations between maternal BMI and offspring DNA methylation were compared to the equivalent estimates for paternal BMI (the negative control), with adjustment for the other parent’s BMI.

Seven cohorts (ALSPAC, CHAMACOS, Generation R, GOYA, MEDALL [INMA and EDEN pooled], NHBCS, RICHS) with the necessary data independently performed robust linear regression to assess the association between paternal BMI (kg/m^2^) and newborn blood DNA methylation at sites identified as associated with maternal BMI. Each cohort ran models adjusted for maternal smoking, age, socioeconomic status, parity and estimated cell counts. We also explored the independent effect of maternal and paternal BMI in mutually adjusted models. Results for each of the seven cohorts were uploaded to the server at the University of Bristol where they were summarised using fixed effects inverse-variance weighted meta-analysis and compared to meta-analysed results of the maternal effect in these seven cohorts. The criteria for evidence of an intrauterine effect were, in the mutually adjusted models, 1) maternal BMI and paternal BMI show the same direction of association with offspring methylation (i.e. paternal BMI is not having an independent effect in the opposite direction to the effect of maternal BMI), 2) the magnitude of association with offspring methylation is larger for maternal BMI than for paternal BMI, 3) there is evidence of heterogeneity (an I^2^ value >40) in a meta-analysis of the maternal and paternal mutually-adjusted estimates. We also calculated heterogeneity P-values between the mutually adjusted maternal and paternal BMI estimates using the metafor R package(96).

### Identification of methylation quantitative trait loci (meQTLs)

We performed a look-up of maternal BMI-associated methylation sites in an online catalogue of both cis- (within 100kb) and trans- methylation quantitative trait loci (meQTLs) identified in an ALSPAC study (http://mqtldb.org/)(35). The meQTLs were identified in cord blood of 771 children at birth using 395,625 methylation probes and 8,074,398 SNP loci after adjustment for sex, the top ten ancestry principal components, bisulfite conversion batch and estimated cell counts. A P-value threshold of 1 *10^-7^ was used to define meQTLs(35). We compared the list of meQTLs to results of an adult BMI GWAS published by the GIANT consortium(36, 58). meQTLs were considered nominally associated with BMI if the GWAS P-value was <0.05. FDR correction for multiple testing was also performed

### Availability of data and materials

Data supporting the results reported in this article can be found in the Supplemental Material (Supplementary File S1). We are unable to make individual level data available due to concerns regarding compromising individual privacy, however full meta-analysis results datasets generated in this study are available from the corresponding author on request.

## Acknowledgements

For all studies, acknowledgements and funding information can be found in the Supplemental Material (Supplementary File S2).

## Authors’ contributions

GCS CR JF SL conceived and designed the study. First authors for each cohort conducted the cohort-specific analyses. GCS, LAS, CM, CA, PY and TME carried out additional analyses for their cohorts. GCS and JF meta-analysed the results. GCS wrote the manuscript with input from all authors. Correspondence and material requests should be addressed to GCS.

